# Daisy quorum drives for the genetic restoration of wild populations

**DOI:** 10.1101/115618

**Authors:** John Min, Charleston Noble, Devora Najjar, Kevin M. Esvelt

**Author notes:** Author's Note on Experimental Pre-Registration: This manuscript is an example of pre-registration to ensure transparency in experimental gene drive research. It's intended as a “living document” that begins by sharing key concepts, rationale, and experimental plans for viewing and comment by the community before any experiments begin. As data are gathered and analyses completed, it will be updated with new figures, and eventually will become one or more peer-reviewed publications. All clinical trials now require pre-registration, and the “registered report” model is gaining traction in psychology, but we're not aware of similar efforts in applied science. This format, which is very much a work in progress, seeks to minimize experimenter effort by making it easy to turn grant proposals into pre-registrations and vice versa while also laying the groundwork for eventual formal publication. Preprint servers offer a way to share the work for external comment while making the document immediately citable in the scientific literature. Openly sharing experimental proposals should accelerate research by allowing scientists to choose whether to collaborate or compete intelligently. For example, if any readers are interested in pursuing these ideas, we would be more than happy to advise, collaborate, or desist in our own efforts as appropriate; we have no desire to duplicate the work of others given the many other urgent projects available. Even if the work is never published in a peer-reviewed journal, the data will remain in place to guide future research along similar lines. It's worth re-emphasizing that pre-registration preprints are readily transformed into grant proposals and publications alike, so the effort of composition is by no means wasted. If peer evaluation of proposals becomes common, this will not only serve to improve experimental designs and increase safety, but could also increase popular or even financial support of the research. In particular, many small-scale philanthropists are interested in backing promising science, but do not have scientific advisors to assess promising proposals; community evaluations of pre-registrations could potentially lead to funding as well as improve the experimental design. Because gene drive systems could alter the shared environment, we believe that all research in the field should be open to ensure that people have a voice in decisions that could affect them. We hope that our colleagues in gene drive research will join us in sharing their experimental plans; however, we understand that scientists also have moral obligations to their students, whose careers are at risk when outsiders “scoop” their best ideas and publish first. We're currently in discussions with scientific journals, funders, policymakers, and intellectual property holders concerning ways to change the scientific incentives governing gene drive research so that researchers can freely share their ideas and plans without fear. If the results from open gene drive research are encouraging, we hope that some future descendant of this pre-registration model will spread to the rest of the experimental sciences. Funding status: The bulk of this work is currently unsupported. We are grateful for a Burroughs Wellcome Fund “Innovations in Regulatory Science” award to study the evolutionary dynamics of gene drive systems in nematode worms, which will help us evaluate daisy quorum once created. Related projects: Daisy-chain gene drive, daisyfield gene drive.

## Abstract

An ideal gene drive system to alter wild populations would 1) exclusively affect organisms within the political boundaries of consenting communities, and 2) be capable of restoring any engineered population to its original genetic state. Here we describe ‘daisy quorum’ drive systems that meet these criteria by combining daisy drive with underdominance. A daisy quorum drive system is predicted to spread through a population until all of its daisy elements have been lost, at which point its fitness becomes frequency-dependent: mostly altered populations become fixed for the desired change, while engineered genes at low frequency are swiftly eliminated by natural selection. The result is an engineered population surrounded by wild-type organisms with limited mixing at the boundary. Releasing large numbers of wild-type organisms or a few bearing a population suppression element can reduce the engineered population below the quorum, triggering elimination of all engineered sequences. In principle, the technology can restore any drive-amenable population carrying engineered genes to wild-type genetics. Daisy quorum systems may enable efficient, community-supported, and genetically reversible ecological engineering.

**Summary:** Local communities should be able to control their own environments without forcing those choices on others. Ideally, each community could reversibly alter local wild organisms in ways that cannot spread beyond their own boundaries, and any engineered population could be restored to its original genetic state. We've invented a 'daisy quorum' drive system that appears to meet these criteria.

“Daisy” refers to a daisy drive, which typically uses a daisy-chain of linked genes to spread a change through a local population while losing links every generation until it stops spreading. “Quorum” reflects the system's ability to “vote” on whether a local population should be altered or not: once all daisy elements are lost, it favors replication by the altered version or the original depending on which is more abundant in the local area. Put together, they result in a change that first spreads through a local population, then either becomes locally prevalent is eliminating, inhibiting mixing at the boundary. All organisms in the target population are altered, but changes are unable to spread much beyond that area due to being greatly outnumbered by wild-type organisms and consequently less able to replicate.

We haven't yet performed any experiments involving daisy quorum systems. Rather, we’re describing what we intend to do, including the safeguards we will use and our assessment of risks, in the hope that others will evaluate our plans and tell us if there's anything wrong that we missed. We hope that all researchers working on gene drive systems - and other technologies that could impact the shared environment - will similarly pre-register their plans. Sharing plans can reduce needless duplication, accelerate progress, and make the proposed work safer for everyone.

## Introduction

Gene drive systems can spread through populations even though they offer no benefit to individual organisms^1–3^. CRISPR-based gene drive systems that use genome editing to copy themselves in place of a target sequence have been described as a potential means of altering diverse sexually reproducing species to benefit public health, conservation, and agriculture^4^. Although this form of gene drive has been demonstrated in four species to date^5–8^, the standard version is self-sustaining, meaning that it is anticipated to spread into every population of the target species around the globe. The possibility of global spread severely limits the utility of gene drive to at most a handful of applications: those aimed at the worst problems, such as malaria and schistosomiasis, and those that would affect only a few countries while clearly posing few ecological risks, such as the elimination of New World screwworm and the alteration of desert locusts to prevent swarming.

Daisy drive systems separate the necessary components of a CRISPR-based gene drive in order to limit the extent of spread to local populations. Daisy-chain drive systems comprise a linked series of elements in which each drives the next in the chain; because the daisy element at the base of the chain does not drive, it is lost in half of offspring, successively depriving subsequent elements of their inheritance advantage^9^. In daisyfield drive systems, many daisy elements all target the same allele and cause it to drive; their number decreases by half with every generation of mating with wild-type organisms until none are left and drive ceases^10^.

By enabling local population editing, daisy drive systems could enable communities to alter their own environments without forcing those choices on others. However, nothing prevents most engineered genes from spreading via normal gene flow to other areas. Ideally, engineered genes would be able to sense the surrounding environment and only persist if there is a local quorum indicating the population should be edited.

Here we describe a simple way to achieve a quorum effect using engineered underdominance and to spread it via daisy drive systems. Daisy quorum drives could help confine alterations within political boundaries and restore any engineered drive-susceptible population, no matter the source, to wild-type genetics.

## Establishing a genetic quorum

The genetic phenomenon of underdominance occurs whenever heterozygotes are less fit than homozygotes (Fig. 1). For example, the offspring of any organism that is heterozygous for a reciprocal chromosomal translocation will inherit two wild-type chromosomes, two rearranged chromosomes, or one of each. Because each translocated arm encodes essential genes – some of these haploinsufficient, meaning that two copies are required for viability – half the offspring of matings with wild-type organisms will be missing an arm, and consequently non-viable. If the two versions are of equal fitness, whichever is more abundant will be more likely to find a mate with the same version, produce more surviving offspring, and eventually go to fixation in the local population.

**Fig. 1.**
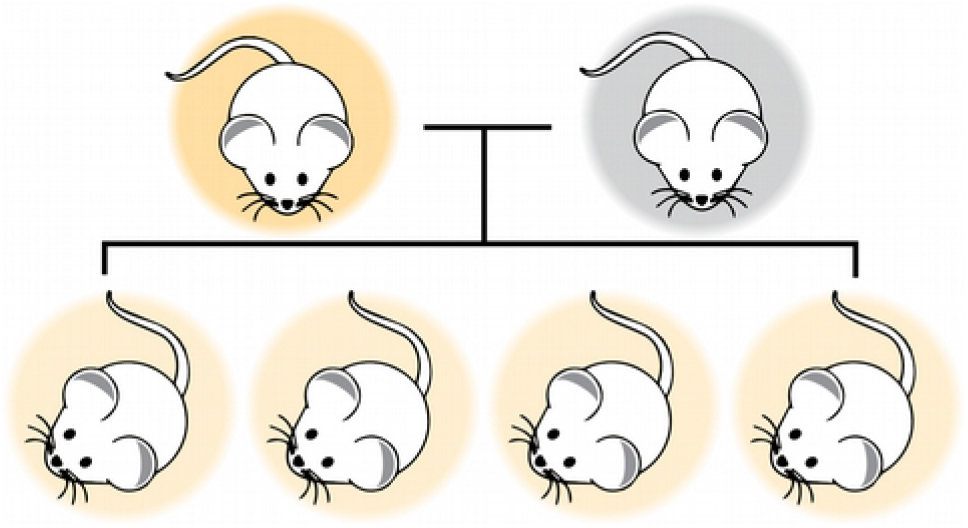
Underdominance and quorum. Two genetic variants exhibit underdominance when hybrids of the two are less likely to reproduce than either parent. The system inhibits gene flow between them and ensures that whichever variant is locally abundant is more likely to find a mate of its own type and produce many descendants. By simply reproducing, the variants determine whether there is a quorum.

Underdominance was the first form of gene drive proposed^11,12^ and the only one to be deployed in the wild^3,13,14^. Since organisms that migrate to another population will be in the minority and consequently are selected against, underdominance approaches with high thresholds pose minimal risk of unintended spread. Because releasing enough organisms of the other type can reduce the local frequency below the threshold, any changes are genetically reversible. These features render underdominance ideally suited for local population alteration, but with two main caveats. First, large numbers of organisms must be released to ensure that the engineered population is in the majority, a constraint that poses particular problems when the target population is an ecologically damaging invasive species. Second, generating healthy strains that display underdominance has historically been extremely difficult. While two toxin-antitoxin underdominance systems have been demonstrated in flies^15,16^, it is unclear how readily these can be adapted to other organisms while maintaining fitness and the extent to which they will be evolutionarily stable. Very recently, a team used site-specific nuclease cleavage to generate reciprocal chromosomal translocations and robust underdominance in flies^17^.

## Linking daisy drive to the quorum effect

We reasoned that daisy drive could enable threshold-dependent elements to locally reach the threshold while releasing a tiny fraction of the organisms that would otherwise be required. Unfortunately, chromosomal translocations occur at low efficiency even when catalyzed by double-strand breaks^17^. While daisy drives could in principle spread toxin-antitoxin systems^15,16^, current approaches to building these may not generalize to other species, nor can *in trans* elements be linked to the quorum effect. Accordingly, we conceived of a way to accomplish a quorum effect using CRISPR alone: by swapping the locations of two haploinsufficient genes (Fig. 2). A daisy drive system can readily replace one gene with another, and because both target genes are essential there is no risk of generating drive-resistant alleles^4,18^. Notably, many haploinsufficient ribosomal genes are under 5kb in length, whereas gene drive cassettes as large as 17kb have been copied with high efficiency^8^. If the target genes are also haploinsufficient in the proliferative germline, any cells that do not correctly copy the drive system will be outcompeted by those that do, potentially eliminating the fitness cost of incorrect repair^9^.

**Fig. 2.**
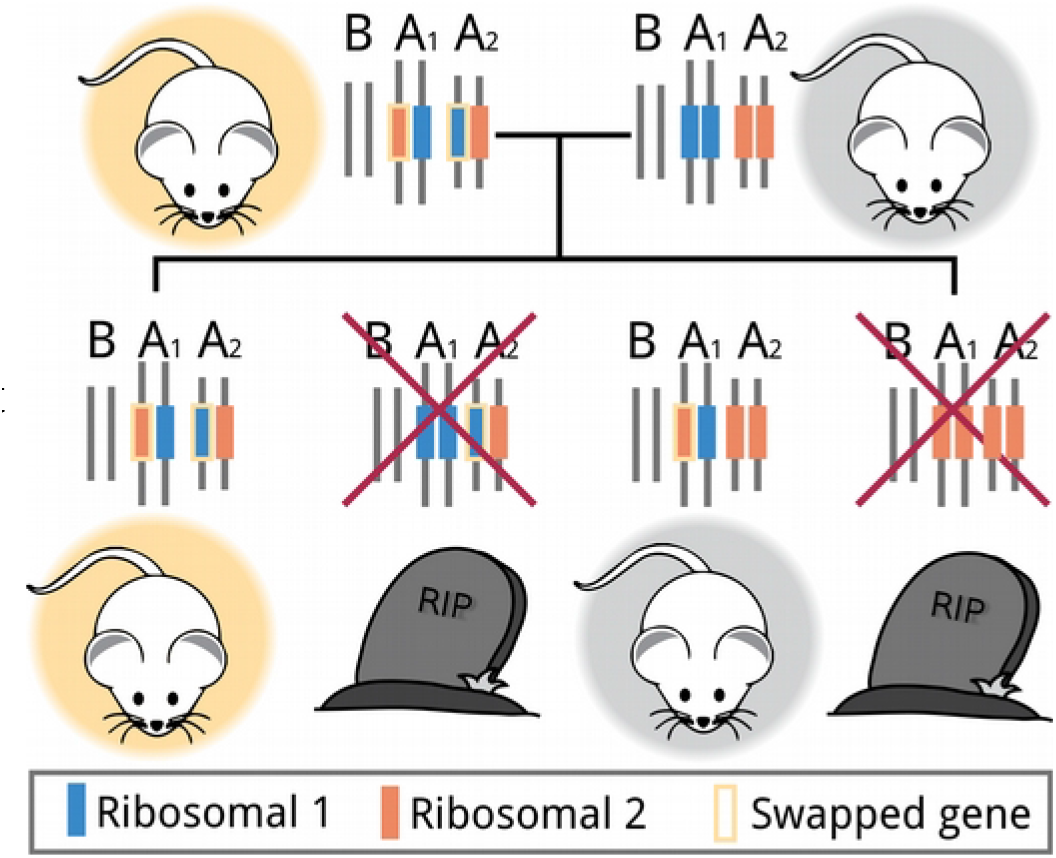
Engineering a quorum. Swapping genes that are haploinsufficient, meaning that two copies are required for viability, results in underdominance relative to organisms with wild-type genetics. When heterozygotes mate with a homozygous swapped or a wild-type organism (pictured), half the offspring fail to inherit two copies of each gene and consequently perish. Many ribosomal genes are haploinsufficient.

In a basic daisy quorum drive system, the daisy elements would target wild-type ribosomal genes A1 and A2, causing them to be replaced with recoded A2* and A1*, plus one or more CRISPR nuclease genes located just downstream of each recoded gene (Fig. 3). Any daisy-chain or daisyfield arrangement can drive both quorum elements simply by targeting A1 and A2. As long as at least one daisy element is present, all offspring will inherit exactly one copy of each recoded ribosomal gene and consequently survive.

**Fig. 3.**
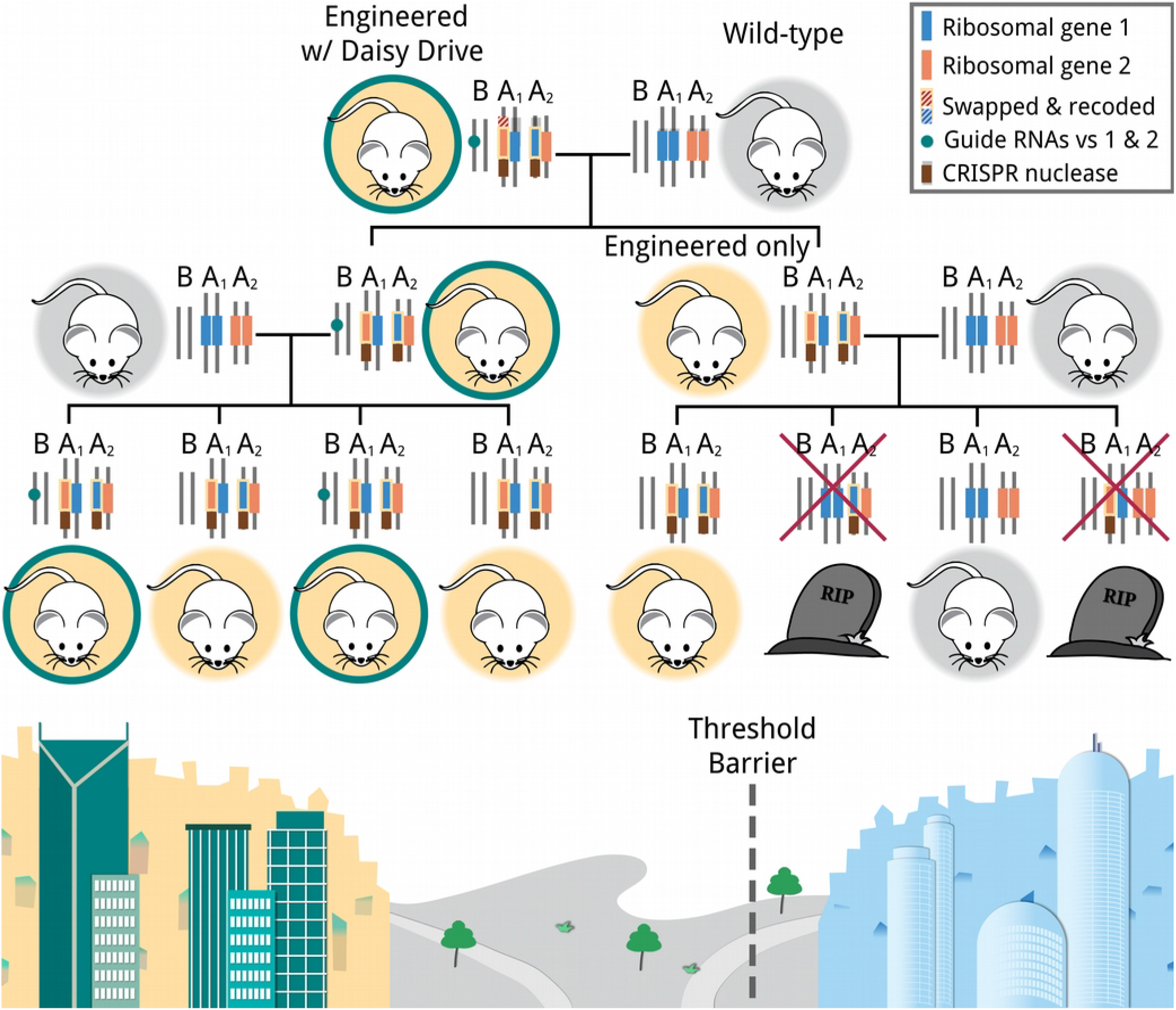
Family tree of a daisy quorum drive system. The animal on the top left has a single intact daisy element remaining, so all offspring of a mating with a wild-type organism inherit one copy of each recoded ribosomal gene and associated CRISPR nuclease. Those offspring that inherit the daisy element (left) display identical dynamics, but those that do not exhibit underdominance (right). Because the quorum effect favors the reproduction of whichever version is in the majority, it is possible to alter a local population in a consenting community (lower left) with very little gene flow to the neighboring community that does not wish to be affected (lower right).

Crucially, every daisy element save for those encoding the nuclease genes must express guide RNAs targeting both wild-type ribosomal genes. These guide RNAs tightly link other elements to the quorum effect by ensuring that any organisms carrying both the guide RNAs and the swapped ribosomal genes with the CRISPR nucleases will only produce gametes that also carry the quorum elements. This will be true as long as CRISPR activity is robust and failure to copy any quorum element is lethal. By the same logic, any genetic cargoes that cannot be located *in cis* to the nucleases might be placed *in trans* if they encode guide RNAs targeting the wild-type ribosomal genes and their own wild-type locus is targeted by the daisy elements in position B. Since there will be at least two CRISPR nuclease genes present, with one or more copy adjacent to each swapped ribosomal gene, CRISPR activity should remain robust for many generations.

## Local and Transient

If released on an island in numbers sufficient for the daisy effect to exceed the quorum threshold, a daisy quorum system will alter the entire island population but spread no further (Fig. 4a). In contrast, a one-time release on a continent will alter the target region, persist for some generations while slowly shrinking, and eventually be eliminated due to the quorum effect and the much larger surrounding population (Fig. 4b). Because all engineered genes are linked to those responsible for the quorum effect, the entire population will eventually return to wild-type unless periodically supplemented with new daisy drive organisms.

**Fig. 4.**
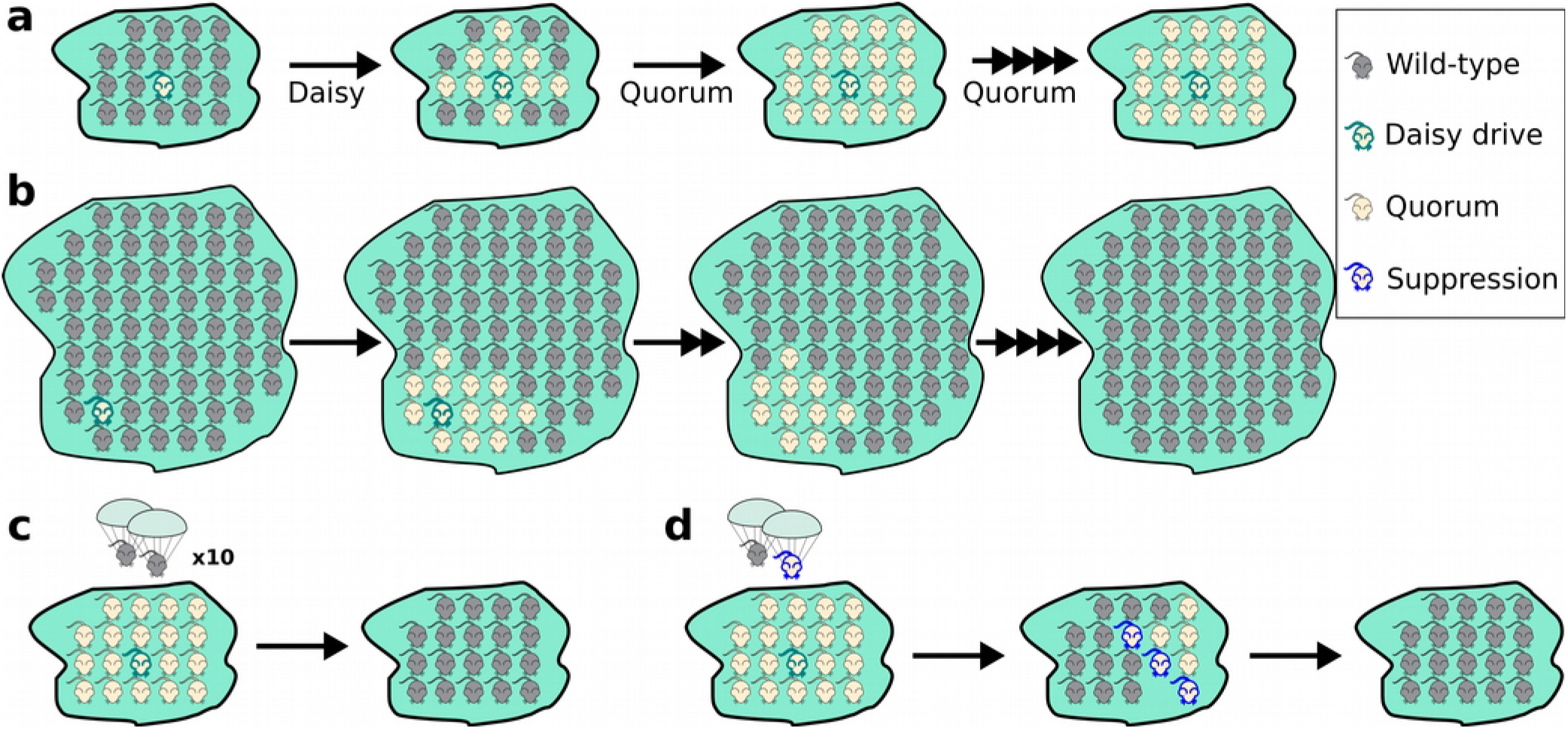
Daisy quorum drive systems enable local, transient, and reversible population editing. **a)** Introduced into a small population, the daisy drive spreads the quorum elements to the majority of individuals. The quorum effect causes it to reach fixation locally, but it cannot invade other populations.**b)** Introduced into a large population, the daisy drive spreads the quorum elements locally. The edited population remains stable for a time if reasonably sized, but is gradually pushed back and eliminated by the greater numbers of wild-type organisms due to the quorum effect. **c)** Introducing enough wild-type organisms to comprise the majority in a population edited by a daisy quorum drive system leads to complete restoration of wild-type genetics. **d)** An easier way to restore wild-type genetics is to release a handful of organisms carrying a suppression element, which spreads infertility through organisms carrying quorum elements, and then wild-type organisms if none are initially present. The quorum effect subsequently eliminates all engineered genes in favor of the wild-type.

## Genetic Reversibility

Populations altered by daisy quorum systems can be returned to wild-type more rapidly by reducing the frequency of the quorum elements below the threshold level. The most direct method is to release sufficient wild-type organisms to exceed the threshold (Fig. 4c). If this would require too many organisms, a far easier approach involves releasing organisms carrying a suppression element that imposes a genetic load exclusively on organisms expressing a CRISPR nuclease (Fig. 4d), plus a suitably diverse population of wild-type organisms if none locally remain. Suppression elements are readily created by replacing a recessive gene required for viability or fertility with a cassette expressing multiple guide RNAs targeting the recessive gene^2,7^. In organisms that carry the CRISPR nuclease, the guide RNAs will cut and replace the target gene in the germline, thereby ensuring that all offspring inherit the non-functional suppression allele. When two such organisms mate, their offspring will be nonviable or infertile, selectively reducing replication of the edited population. Once below the threshold, the quorum effect will eliminate every remaining copy.

## Daisy restoration drive: completely removing unwanted global drive systems

Global gene drive systems are theoretically capable of spreading to every population of the target species in the world. Neither pure reversal drives carrying only guide RNAs nor gene drive 'brakes'^19^ can eliminate every copy of an unwanted global drive system, rendering them inadequate defenses against accidents or misuse. While the phenotypic effects of a rogue drive can be countered by releasing another drive system to overwrite it and immunize unaffected wild populations^4^, such immunizing reversal drives are themselves global and will leave residual CRISPR components in every population of the species.

However, any preexisting daisy drive system can be adapted to overwrite all copies of an unwanted global drive system that uses a different CRISPR nuclease. Replace the latter with a cassette encoding two types of guide RNAs. The first set uses the unwanted drive nuclease to cut target sites present in the unwanted system that it does not share, and also drives all daisy elements (Fig. 5a, depicted for daisy quorum). The second set uses the daisy nuclease to cut the wild-type locus, thereby immunizing local organisms affected by the daisy drive system. When heterozygous with the unwanted drive, the daisy system will avoid being cut and drive all daisy elements; it will act like a global immunizing reversal drive only in the presence of its target. When heterozygous with wild-type alleles, it will behave as a normal daisy drive. This combination of traits allows it to spread efficiently through an already-edited population, then spread through the wild-type organisms enough to eliminate every copy of its target. While any daisy drive system can be adapted to locally eliminate every copy of a rogue global drive system without affecting distant populations, the addition of quorum enables the population to be restored to wild-type genetics afterwards (Fig. 5b). We refer to this combination as a daisy restoration drive.

**Fig. 5.**
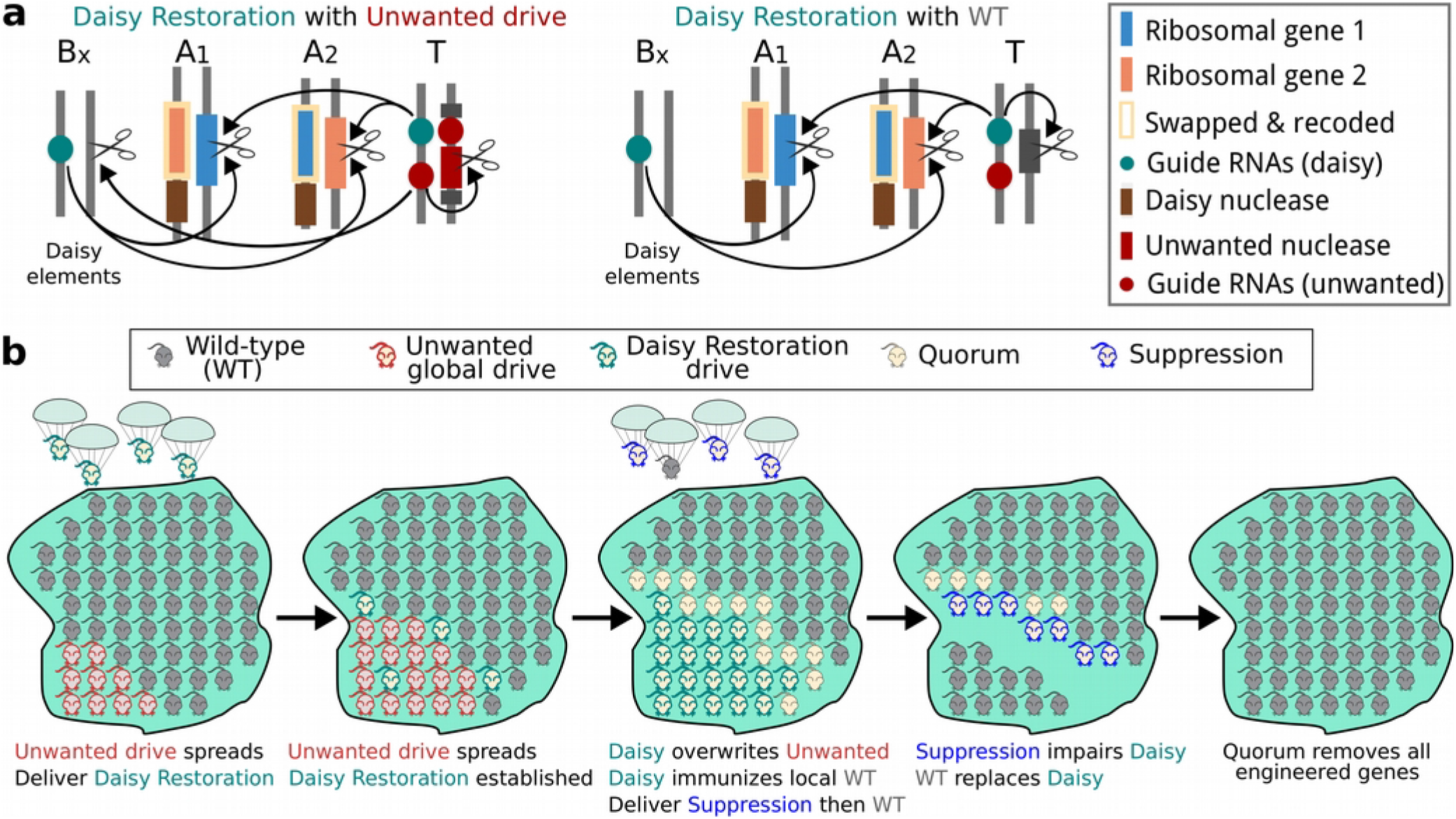
Daisy restoration drives can restore populations altered with global drives to wild-type. **a)** An unwanted global drive system can be countered by an appropriate daisy restoration drive, which is a daisy quorum drive system with an immunizing reversal element. This element has guide RNAs that use the unwanted drive nuclease to spread itself and all daisy elements, thereby acting like a global drive, and can also immunize local wild-type organisms using the daisy nuclease and guide RNAs encoded by the penultimate daisy element. **b)** Suppression and quorum lead to restoration of wild-type genetics.

## Daisy restoration: returning any engineered population to wild-type genetics

In theory, daisy quorum drive systems can overwrite *any* engineered change in a target population, then restore wild-type genetics. To remove an unwanted sequence from a population, insert in its place a cassette encoding guide RNAs that target the unwanted sequence as well as the ribosomal genes used for the quorum system, and optionally the other daisy elements (Fig. 6a). Every copy of the target sequence will be replaced by the guide RNA cassette, which is tightly linked to the quorum effect (Fig. 6b). Subsequently eliminating the quorum elements should remove every last engineered gene, another form of daisy restoration drive.

**Fig. 6.**
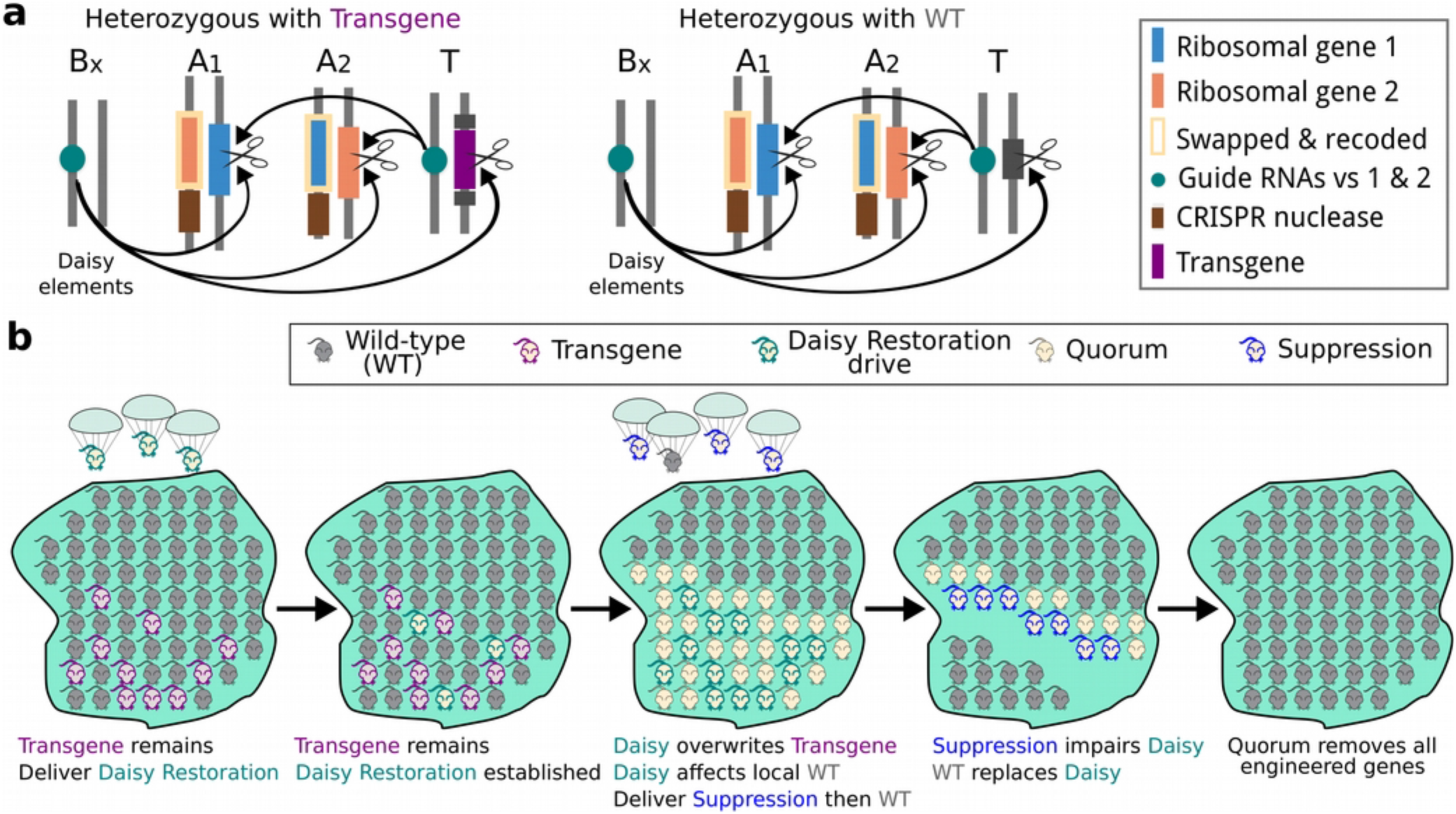
Daisy restoration drives can restore arbitrary engineered populations to wild-type genetics. **a)** Daisy restoration drives can precisely remove any target sequence, including any engineered gene, from local populations. This type of daisy restoration drive is created by combining a daisy quorum system with an overwriting element inserted in place of the target sequence that encodes guide RNAs that drive all daisy elements. One of the daisy elements must also be adapted to cut the target sequence, thereby driving the overwriting element when the target is present. **b)** Once all copies of the target sequence have been overwritten, suppression and quorum restore the population to wild-type genetics.

## Engineering challenges and considerations

The primary obstacle to constructing a daisy quorum system is the fact that the positions of haploinsufficient genes cannot be exchanged sequentially without causing lethality. Since the efficiency of CRISPR-mediated insertion is low, achieving simultaneous quadruple replacement of all four copies is extremely unlikely. Instead, we suggest replacing each target ribosomal gene with a recoded version of itself that is flanked by homology regions matching the ends of the other gene as well as by irreversible recombination sites (Fig. 7). Expressing the recombinase will swap the positions of both pairs of recoded genes in a single reaction. Cargo genes and germline-expressed CRISPR nuclease genes can be added to each of the copies via CRISPR or by recombinase-mediated cassette exchange, then combined with a daisy drive system whose penultimate element targets both wild-type haploinsufficient genes.

**Fig. 7.**
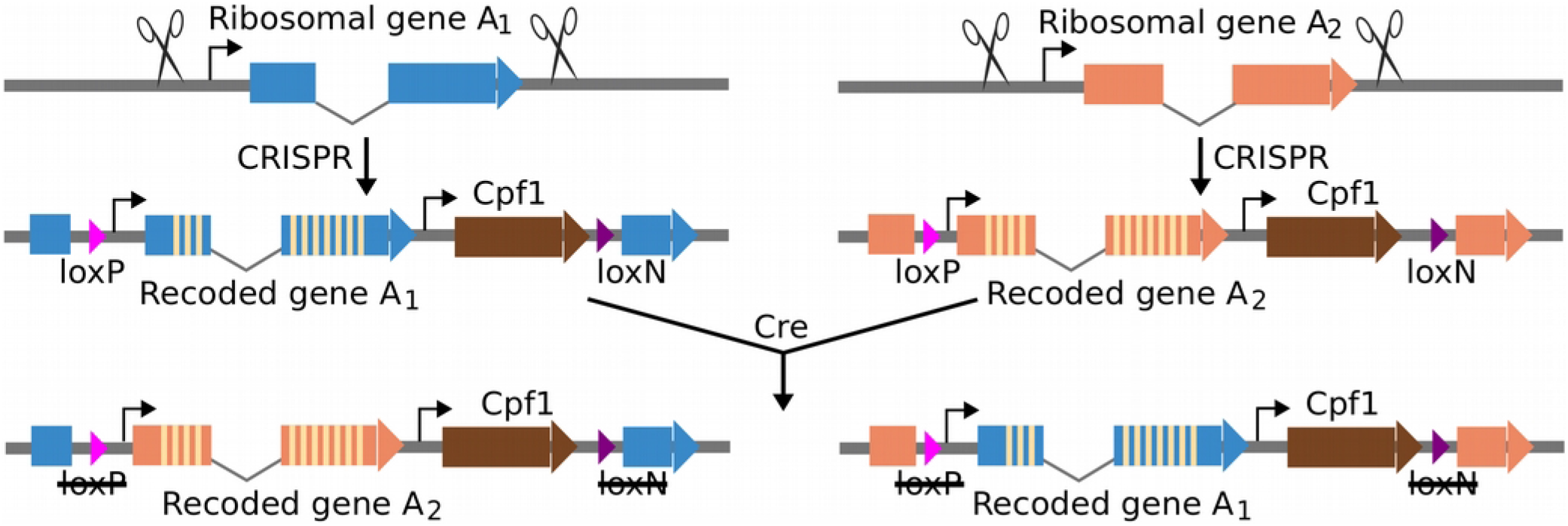
Ribosomal gene rearrangement. We intend to build daisy quorum drives by using CRISPR to make cuts on both the 5' and 3' ends of endogenous ribosomal genes. A recoded version of the same gene is then inserted via homology-directed repair with flanking lox sites and homology to the 5' and 3' ends of the gene. The resulting organisms are then crossed and the recoded ribosomal genes swapped via Crerecombinase.

There are arguably three major potential barriers to implementing efficient daisy quorum drive. First, one might be concerned that daisy quorum and particularly daisy restoration simply involve too many moving parts to work reliably, especially on an population level. Were the designs based on any other biotechnology, this would indeed be prohibitive. However, daisy quorum systems rely entirely on CRISPR. Published experiments routinely report cutting rates of 100% even using just one guide RNA^6–8,20^, let alone several, so cutting is known to be sufficiently reliable^21^. Second, it may not be possible to target so many sites at once without risking second-order effects. Yet multiplex cutting is becoming increasingly feasible in multiple species^22,23^, and while there are not currently enough sequence-divergent guide RNAs available to build a powerful daisy restoration drive system without risking recombination, extending our earlier approach to identifying such guide RNAs^9^, particularly for CRISPR nucleases such as Cpfi^24^, will soon yield more than enough.

Third, recoding ribosomal genes may result in unacceptably high fitness costs. However, it is worth noting that because the genes will swap positions, only the PAM of each target site need be edited; almost all of the coding sequence will remain intact. More extensive recoding efforts in bacteria^5,26^ and fruit flies^16^ suggests that problems are unlikely to occur and surmountable in organisms amenable to more than one design-build-test cycle.

Most construction criteria for daisy quorum systems are the same as for daisy-chain and daisyfield drive systems. Daisy elements encoding guide RNAs can be located anywhere euchromatic, but are subject to the same stability considerations as global drive systems. That is, the cargo element and the CRISPR nuclease gene should be located adjacent to the recoded version of a conserved gene whose disruption severely impairs replication, or within a gene whose disruption is the goal; the most parsimonious solution is to include them adjacent to the swapped haploinsufficient genes. The wild-type version of the targeted gene must be cut with multiple guide RNAs in order to ensure that cutting will occur regardless of sequence diversity and that repair by any method other than copying the drive element is at least as costly to the organism as is the drive system. This precaution is necessary to ensure that there are no alleles that cannot be cut, because these might otherwise outcompete the cargo. Since the other daisy elements are not intended to reach fixation anyway, drive-resistant alleles can only reduce the efficiency.

For safety reasons, no daisy drive component may directly or indirectly target both its own wild-type allele and any allele corresponding to a CRISPR nuclease gene. Any such system would comprise a multi-element global drive, with the nuclease and guide RNAs together driving the elements encoding them^9^. Sequence similarity between elements carrying different guide RNAs must be minimized to prevent possible recombination events. Studies of stability in extremely large populations of species chosen for the purpose will be necessary to measure the risk of recombination as a function of the population size to be altered. However, it is worth noting that any such global drive system could be also countered and the population restored to wild-type using the same daisy quorum approach as for a rogue drive system.

Finally, it's worth noting that daisy quorum systems afford a potential way to improve the safety of laboratory studies involving global gene drive systems. Despite calls for regulation and recommendations of laboratory safeguards even before the technology was demonstrated, very few countries have updated their policies accordingly, and few other levers can enforce compliance. Since the primary risk posed by rogue drive systems is a social backlash against biotechnology in general and gene drive in particular, a technological means of restoring populations to their original genetics could be a powerful shield against the consequences of accidents and misuse. It may be wise to publicly construct daisy quorum systems in most species that might be subjected to accidental or deliberate rogue drive releases, thereby enabling their swift restoration to wild-type genetics.

## Discussion

By providing a means of restoring arbitrary engineered populations to their original genetics, daisy quorum systems offer a morally necessary safeguard against real and perceived damage to the shared environment. Real damage refers to genetic changes with unwanted ecological consequences. Many, perhaps most changes will have no such consequences, but are nonetheless a source of profound concern.

The spread of engineered genes into wild populations is deeply distressing to people who revere the perceived wilderness and do not wish it sullied by any form of human intervention. Critics rightly point out that humanity has been engineering domesticated plants and animals using selective breeding for millennia, that gene flow between species is common throughout the natural world, that horrific disease and suffering are endemic in the wild, and that technology and all its benefits are by definition unnatural. All of these are immaterial. Many people are profoundly disturbed by things they *perceive* to be unnatural, and their suffering at the perceived damage to the object of their reverence is genuine. While society cannot and should not abandon the benefits of engineered organisms, we are obligated to do what we can to abide by their wishes and minimize the unwanted alteration of wild populations.

Daisy quorum systems offer a genetically reversible way to make agreed-upon changes and keep them confined to consenting polities. Just as important, they are theoretically capable of restoring any drive-amenable population to its original genetic state.

The anticipated development of readily accessible global CRISPR drive systems capable of unilaterally altering a target species over time poses severe challenges for the democratic governance of shared ecosystems. Even if a global drive system does not affect every individual organism, it is likely to eventually affect every population of the target species in the world, if only due to unauthorized transport by humans. While the release of a rogue drive system is unlikely over the next decade or so, the increasing availability of genome editing technology renders it inevitable. Should anyone accidentally or unilaterally set in motion a process that would ultimately edit the bulk of an entire wild species, the damage to public trust in scientists and governance would be severe and long-lasting.

By offering a way to restore populations to their original genetic state, daisy quorum drives may become a critical tool for preserving natural ecosystems and public confidence in biotechnology.

## Experimental plans (pre-registration)

We intend to build and test daisy quorum drive systems in the nematode worm *C. brenneri.* Ribosomal genes targeted for swapping include rpS6, rpS12, rpS20, and rpL14. We are currently working to assemble a more accurate *C. brenneri* genome in order to verify that these genes are on different chromosomes. Using *in vitro* purified Cas9 and guide RNA complexes, each ribosomal gene will initially be replaced with a version of itself in which selected internal target sites for Cpf1 and Cas9 have been eliminated (Fig. 7). Genes encoding Cpf1, Cas9, or both nucleases will be inserted downstream. All daisy quorum elements in nematodes will additionally carry a fluorescent protein marker such as mCherry to enable easy tracking. The daisy quorum components will be delivered to the nematodes via computer-assisted microinjection of live worms using our custom-built apparatus.

When simulating restoration of populations affected by an unwanted drive system, the target “unwanted drive” will consist of a Cas9 gene (for a daisy quorum drive based on Cpf1; else the two will be switched) and guide RNA cassette targeting GFP. It will be crossed with a GFP background “wild-type” strain. Spread of the unwanted drive can be monitored through the loss of GFP signal and gain of a BFP marker associated with the guide RNA.

Most experiments will be performed in cultures of nematodes on tissue-culture plates. This format allows for establishing multiple discrete populations on the order of tens of thousands to millions depending on well size. Gene flow between the populations can be simulated by mixing the individual wells via pipetting. Real-time tracking of gene drive spread will be accomplished using a plate-reader. Studies of the evolutionary stability will be conducted in serially linked flask populations containing hundreds of millions of nematodes each, in which a small culture volume is transferred between adjacent flasks in each generation. The initial number of daisy drive worms will be calculated to preclude spread to all populations; if this event occurs, it is a sign of an unwanted recombination event.

If successful, we may eventually consider building daisy quorum drive systems in other organisms, at which point we will update this document with our new plans.

## Safeguards

As *C. brenneri* is a tropical species that is not found in the New England region, all experiments will by default employ ecological containment. Molecular confinement in the form of split drive, synthetic site targeting, or both will be implemented at all times. These safeguards consequently meet or exceed the published safety guidelines for laboratory research involving gene drive systems^27^. All experiments will be performed and nematode stocks kept in secured laboratory environments with controlled access.

## Acknowledgements

We thank George Church, Luke Alphey, Floyd Reed, Alun Lloyd, Fred Gould and Bruce Hay for helpful conversations over the past several years that refined our thinking on underdominance.

